# Characterising the vocal repertoire of the Indian wolf (Canis lupus pallipes)

**DOI:** 10.1101/612507

**Authors:** Sougata Sadhukhan, Lauren Hennelly, Bilal Habib

## Abstract

Vocal communication in social animals plays a crucial role in mate choice, maintaining social structure, and foraging strategy. The Indian grey wolf, among the less studied subspecies, is a social carnivore that lives in groups called packs and has many types of vocal communication. In this study, we characterise harmonic vocalisation types in the Indian wolf using howl survey responses and opportunistic recordings from captive and nine packs (each pack contains 2-9 individuals) of free-ranging Indian wolves. Using principal component analysis, hierarchical clustering, and discriminant function analysis, we found four vocal types using 270 recorded vocalisations (Average Silhouette width Si = 0.598) which include howls and howl-bark (N=238), whimper (N=2), social squeak (N=28), and whine (N=2). Although having a smaller body size, Indian wolf howls have an average mean fundamental frequency of 0.422KHz (±0.126), which is similar to other Holarctic clade subspecies. The whimper showed the highest frequency modulation (37.296±4.601 KHz) and the highest mean fundamental frequency (1.708±0.524 KHz) compared to other call types. Less information is available on the third vocalisation type, i.e. ‘Social squeak’ or ‘talking’ (Mean fundamental frequency =0.461±0.083 KHz), which is highly variable (coefficient of frequency variation = 18.778±3.587 KHz). Our study’s characterisation of the Indian wolf’s harmonic vocal repertoire provides a first step in understanding the function and contextual use of vocalisations in this social mammal.

## Introduction

Vocalisation plays a critical role in social animals for conveying information on foraging, reproductive, and social behaviours [1–7]. Characterising the vocal repertoire of a species provides a base for understanding the behavioural significance of different vocalisations and studying how vocal communication varies across the populations, subspecies, and taxa [8–10]. The wolf (*Canis lupus*) is a social mammal and uses a variety of vocalisations for communication. Being present throughout Eurasia and North America, the wolf is one of the most widely distributed land mammals and occupies a wide range of different habitat types [11]. The Indian wolf is among the smallest [12] and one of the most evolutionarily distinct wolf subspecies, having diverged around 270,000 and 400,000 years ago based on mitochondrial DNA [13–15]. Studying the vocal repertoire of the Indian wolf can aid in future studies on the function of different vocal signals in Indian wolves and, more broadly, the variation in vocalisation across subspecies and taxa within the *Canis* clade.

The best-known wolf vocalisation – the howl – is a long-range harmonic call used for territorial advertising and social cohesion [1,16–18]. Recent studies have shown 21 different howl types across various canid subspecies based on quantitative similarity in modulation pattern [8]. Along with howl, wolves also communicate using 7 to 12 other harmonic calls, which is a clear pitch sound wave that possesses multiple integral frequencies [19–21]. Many of these other harmonic vocalizations are short-ranged, and due to difficulties in recording these calls, remain less studied compared to the wolf howl [22]. These short-ranged calls are important for communicating passive or aggressive behaviour among social canids [22–24].

The whimper, whine and yelp are various calls for communicating passive and friendly behaviour among wolves [18,23], whereas noisy calls, which don’t have a clear pitch or distinct frequency band in their spectrograms, communicate different levels of aggression [18,23]. The whimper and whine vocalization is similar to a crying sound with the whimper having a comparatively shorter duration than whine [18,25]. The whine vocalization is mostly used for submissive behaviour whereas the whimper is primarily used for greeting [18]. Yelp is a short and sharp cry and is used in submissive behaviour with body contacts [18,25]. To communicate different levels of aggression behaviors, wolves use noisy calls which consist of the growl, woof, and bark. Growl is a laryngeal sound to show dominance in any interaction, whereas the woof vocalization is a non-vocal sound cue (without involvement of vocal cords) used by adults for their pups [18,25,26]. The bark is a short low pitched sound with rapid frequency modulation and is used during aggressive defence [19,26], such as defending pups or defending a food resource. Wolves also express communication through mixed vocalisation either by ‘successive emission’ or by ‘superimposition’ of two or more sound types [21]. A recent study on Italian wolf (*Canis lupus italicus*) suggests six other types of calls may combine with howls to make a complex chorus vocalisation [27].

This study investigates the acoustic structure of harmonic vocalizations of Indian wolves and classifies harmonic vocalizations using a statistical approach. We accumulated the vocalisation data from free-ranging and captive Indian wolves, which will be the first study to evaluate different types of vocalisations of this wolf subspecies. Using multivariate analyses, we describe and classify different harmonic calls to develop a vocal repertoire of the Indian wolf.

## Materials and methods

### Study Species

The Indian wolf (*Canis lupus pallipes*) is among the smallest subspecies of wolf with an average body weight is 20.75 kg [12]. Indian wolves are mostly found in grasslands and the edges of dense forest on the Indian subcontinent [12,28–31]. We recorded vocalisations from nine packs of free-ranging wolves and ten captive wolves from Jaipur Zoo. For captive wolves, we collected vocalisation data from 10 wolves: two adult pairs and six subadults. One adult male was recently captured from wild near the city of Jaipur, Rajasthan, India. The rest of the Indian wolves are descendents of captive breeders at Jaipur Zoo.

### Study Sites

The study was conducted in the state of Maharashtra (Figure 1) and Jaipur Zoo of Jaipur, Rajasthan, India. The study site of Maharashtra includes the semi-arid area of central Deccan Plateau [32], which consists overlapping habitat of Tropical Dry Deciduous Forest, Grassland, Savanna (Western Part) and Tropical Moist Deciduous Forest (Eastern Part) [33].

**Figure 1.**
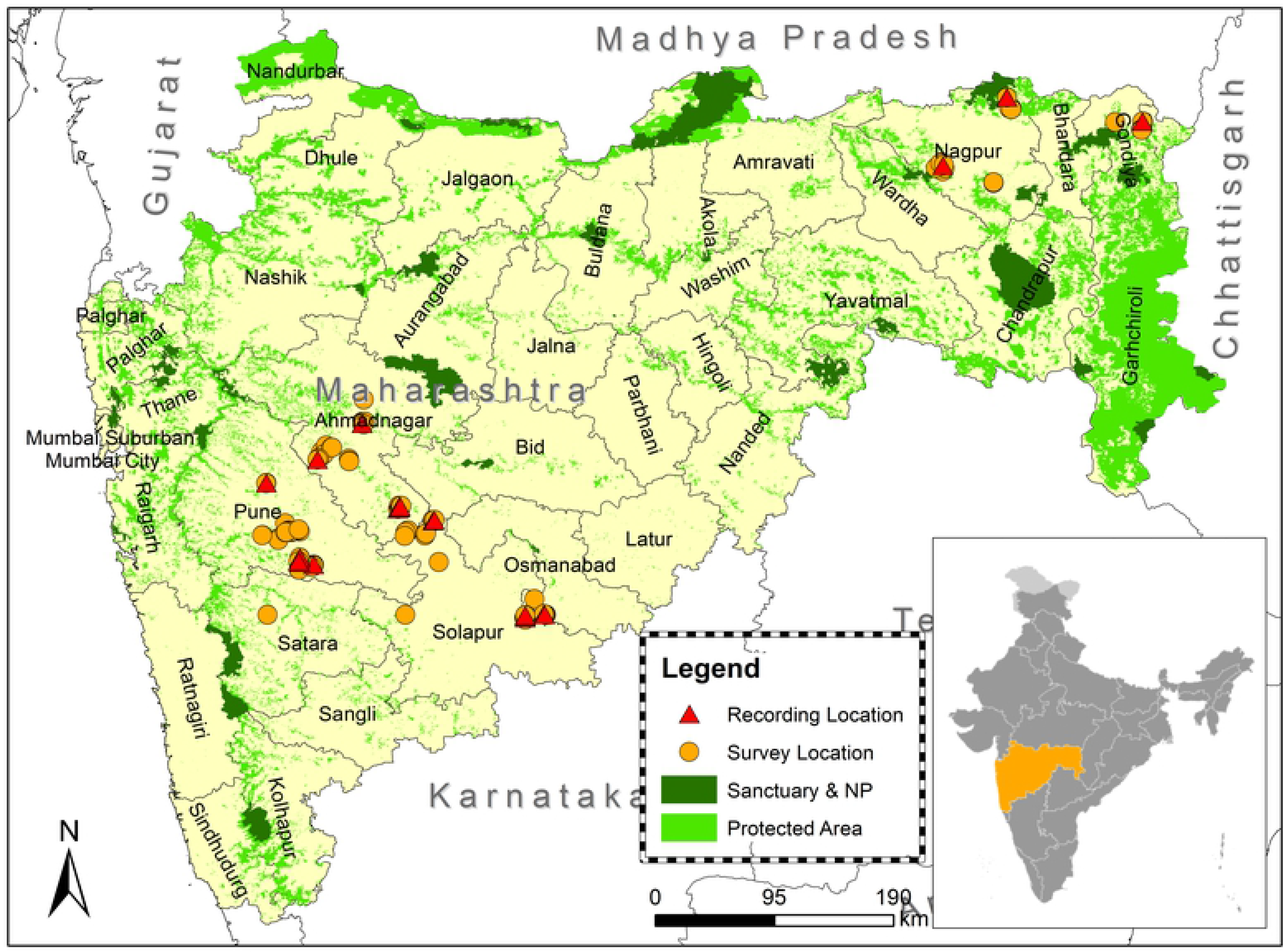
Map of survey sites of the free-ranging wolves.

**Figure 1.**
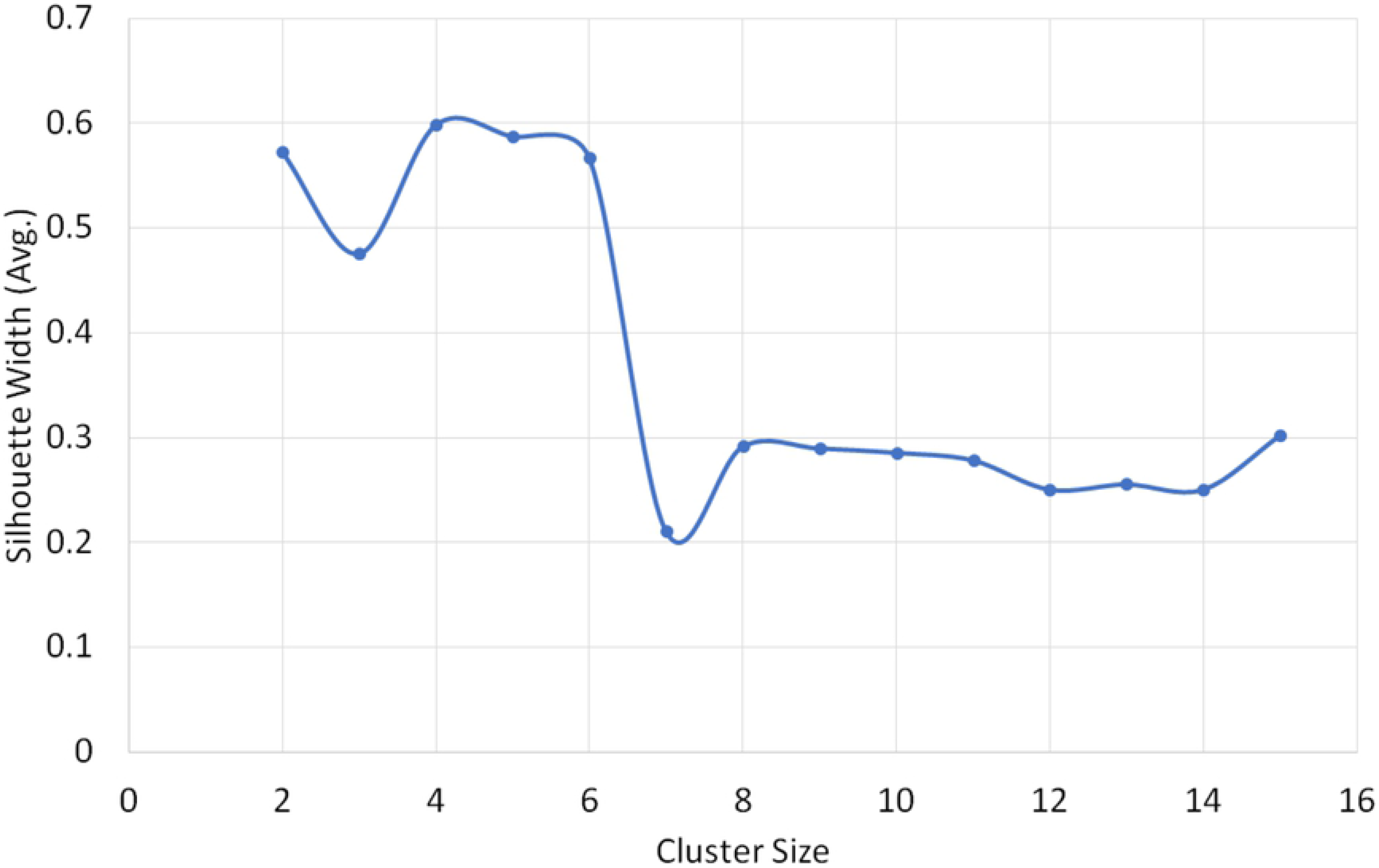
Average silhouette width plotted against 14 different solutions (2-15 cluster). Average Silhouette width represents the significance level (0 represents poor fit, 1represents best fit). We obtained maximum silhouette width in 4 cluster solutions i.e. S_i_= 0.598.

### Data Collection

Vocalization of free-ranging wolves was recorded through acoustics survey from November 2015 to June 2016. Most of the long-distance vocalisation data were collected through howling surveys to elicit howl behaviour. For other types of vocalisation data, we relied on opportunistic recordings from free-ranging wolves and captive wolves, in which installed microphones near the enclosure was used to record vocalizations. Howl surveys were performed during early morning and evening hours using pre-recorded howls that were previously recorded from the Jaipur Zoo Indian wolves. Each howling session consisted of five trials with three-minute intervals. A 50-second-long pre-recorded solo howl was played thrice using JBL Xtreme speakers [34] in the order of increasing volume. The session was followed by two 50-second-long chorus howls with an increase in volume level. In the case of howling response, the session was terminated and repeated after 15 to 20 minutes. Responses were recorded using Blue Yeti Pro Microphone [35] attached with Zoom H4N Handheld Audio Recorder [36] at a sampling rate of 44.1KHz on 16-bit depth with 80 Hz noise filter. Along with survey responses, the opportunistic recording sessions were conducted near Indian wolf den sites and rendezvous sites. In addition to howl surveys at Jaipur Zoo, vocalisations of captive wolves were recorded by installing microphones in the front of cages during closing hours (6:30 pm-7: 30 am).

### Feature Extraction

Spectrogram of each vocalisation was generated through the Raven Pro 1.5 software [37] using the *Discrete Fourier Transform* (DFT) algorithm. The discrete Fourier function transforms same length sequence of equally spaced sample points (N), (N is a prime number) with circular convolution being implemented on the points [38]. *Hann windows* were used at the rate of 1800 samples on 35.2 Hz 3dB filter. A total of 270 spectrograms were selected for further analysis based on clarity (i.e. clearer spectrogram with low noise and without external sound overlap). Web plot digitiser v3 [39] was used to digitise fundamental frequency from the spectral images. This digitised data was obtained at 0.1sec resolution. From this data, eleven acoustic variables (Table 1) were obtained based on their performance from previous studies [40,41].

**Table 1.** Acoustic variables based on fundamental frequency (f_0_) that were extracted for this study

| Variable Name | Definition of Variable |
| --- | --- |
| Min f | The minimum frequency of the fundamental ($f_0$) |
| Max f | The maximum frequency of $f_0$ |
| Range f | Range of $f_0$; $f_0 = \text{Max } f - \text{Min } f$ |
| Mean f | Mean frequency of $f_0$ at 0.1 s interval over the duration |
| Duration | Duration of Howl measured at $f_0$; Duration = $t_{\text{end}} - t_{\text{start}}$ |
| Abrupt<sub>0.025</sub> | Number of abrupt changes in $f_0$ more than 25Hz at single time step (0.1sec) |
| Abrupt<sub>0.05</sub> | Number of abrupt changes in $f_0$ more than 50Hz at single time step (0.1sec) |
| Abrupt<sub>0.1</sub> | Number of abrupt changes in $f_0$ more than 100Hz at single time step (0.1sec) |
| Stdv | Standard Deviation of $f_0$. |
| Co-fm | Coefficient of frequency modulation of $f_0 = \Sigma \vert f(t) - f(t+1)\vert / (n-1) \times 100 / \text{Mean } f_0$ |
| Co-fv | Coefficient of frequency variation of $f_0 = (SD/\text{mean}) \times 100$ |

### Statistical Analysis

#### Principal Component Analysis (PCA)

To obtain a smaller set of representative variables, we used principal component analysis (PCA), which extracts linearly uncorrelated variables from a suite of potentially correlated variables. PCA is advantageous to building a robust model more quickly than would be possible with raw input fields [42]. To simplify the interpretation of factors, we performed varimax rotation using Kaiser normalisation. From our dataset of 270 vocalisations, we used eight scalar variables that are related to spectral structure (Range f, Duration, Abrupt0.025, Abrupt0.05, Abrupt0.1, Stdv, Co-fm, Co-fv) for PCA analysis through the software SPSS (v22). The first principal component (PC1) and second principal component (PC2) were used in the subsequent clustering analyses.

#### Cluster Analysis

To classify the recorded vocalisations from the Indian wolf, we used agglomerative hierarchical clustering through the R package AGglomerative NESting (AGNES). The agglomerative hierarchical clustering algorithm measures the dissimilarity between single and groups of observations using a “bottom-up” approach, thereby constructing clusters [43]. Agglomerative hierarchical clustering was performed using Euclidean distances with PC1 and PC2 from the 270-vocalisation data using eight scalar variables. Subsequently, s*ilhouette clustering* was combined with AGNES to validate the number of clusters in our vocalisation data. Silhouette clustering measures the similarity of observation within its cluster compared to other clusters [44]. The average silhouette value (0 represents poor fit, 1 depicts the highest fit) describes *the evaluation of clustering validity* [44]. Average Silhouette width (S_*i*_) was calculated for 14 different solutions (2 to 15 clusters). The “solution” that provided the best fit was selected upon the maximum average silhouette value. The dendrogram was plotted using ‘*Circlize Dendrogram*’ in the package ‘*Dendextend’* in program R.

#### Discriminant Function Analysis (DFA)

Discriminant function analysis (DFA) was performed using PC1 and PC2 as an independent variable under the program SPSS v22 to cross-validate the obtained clusters from AGNES. Predicted clusters that were determined by the maximum silhouette value were then used as a grouping variable to see within-group covariance in DFA analysis.

#### Box plot using simple scalar variable

From these clusters, we then used box plot to show the overall pattern and distribution characteristics of different vocal clusters.

## Results

### Principal Component Analysis

Two principal components (PC1 and PC2) were generated from the 8-simple scalar variables through PCA based on Kaiser-Guttman Rule (Eigenvalue >1) [45]. PC1 and PC2 together explained 70.6% variance. PC1 was based on the variances of six acoustic parameters (Abrupt_0.025_, Abrupt_0.1_, Abrupt_0.05_, Co-fv, Range f, Stdv) whereas PC2 is explained by the variances of five parameters (Abrupt_0.1_, Co-fm, Duration, Range f, Stdv) (Table 2)

**Table 2.** The loadings of PC1 and PC2. Communalities are the proportional factors by which the importance of each variable is explained.

| Acoustics Parameters | PC1 Loading after varimax rotation | PC2 loading after Varimax Rotation | Communalities |
| --- | --- | --- | --- |
| Abrupt<sub>0.025</sub> | .858 | 0 | 0.741 |
| Abrupt<sub>0.1</sub> | .602 | .415 | 0.535 |
| Abrupt<sub>0.05</sub> | .825 | 0 | 0.732 |
| Co-fm | 0 | .945 | 0.898 |
| Co-fv | .862 | 0 | 0.750 |
| Duration | 0 | -.382 | 0.231 |
| Range f | .852 | .392 | 0.880 |
| Stdv | .759 | .556 | 0.885 |
| % of Variance | 48.946 | 21.692 |  |

### Cluster Analysis

The highest silhouette value (S_*i*_ = 0.598) was obtained at the 4-group solution in the cluster analysis using PC1 and PC2 from PCA analysis (Figure 2). The average silhouette value was 0.62 for the first cluster (N=238), 0.37 for the second cluster (N=2), 0.38 for the third cluster (N=28) and 0.73 for the fourth cluster (N=2) (Figure 3). The 4 clusters were formed at 3.9 clustering scale through agglomerative hierarchical clustering (Figure 4).

**Figure 2.**
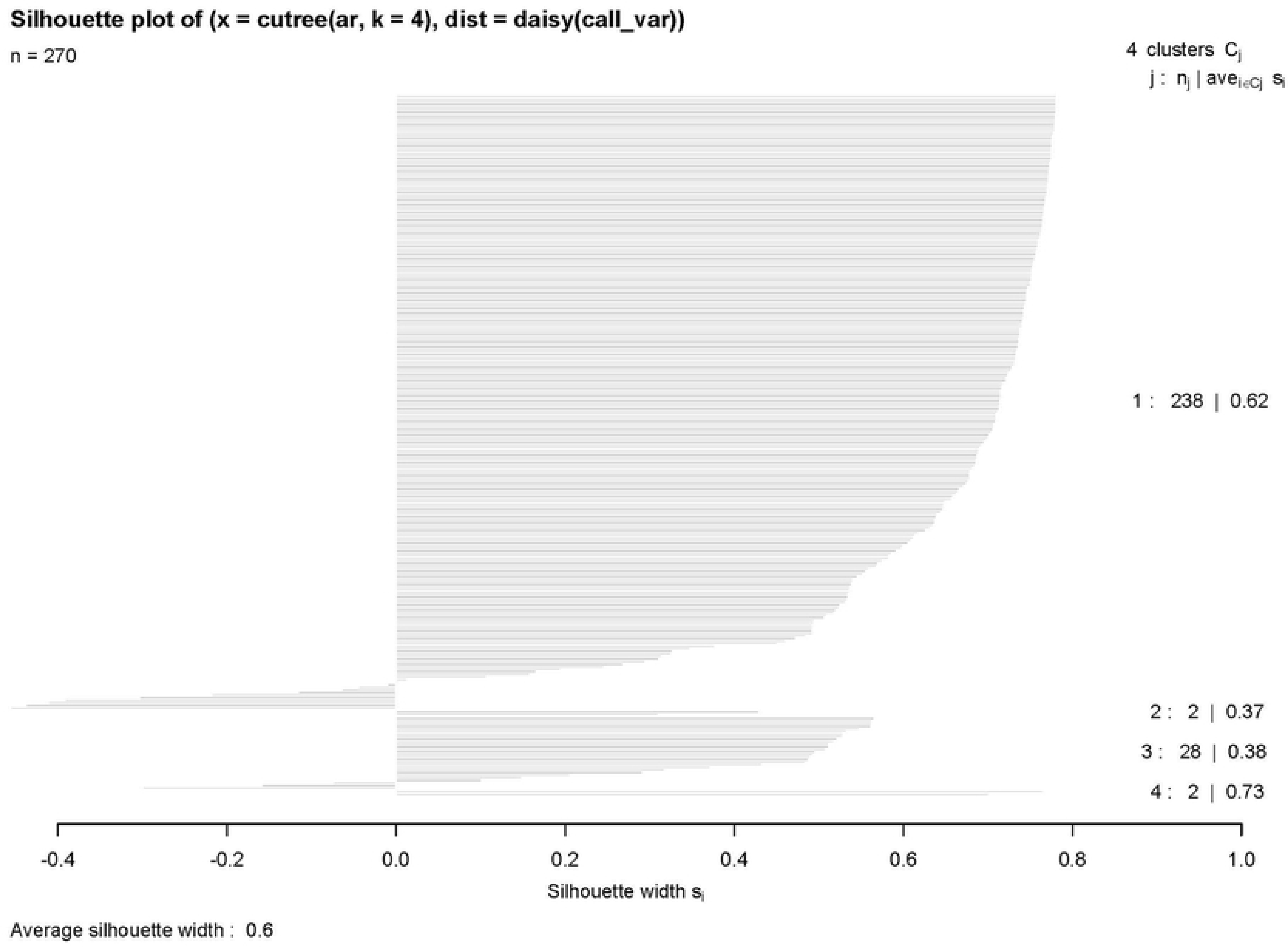
Silhouette plot is a validation method for the consistency among the clusters. It provides information about the similarity or difference of each call from its clusters. A negative value indicates the chance of a call to fall under the neighbouring cluster. The average silhouette value of 4 groups are 0.62 (N=238), 0.37 (N=2), 0.38 (N=28), 0.73 (N=2) respectively.

**Figure 3.**
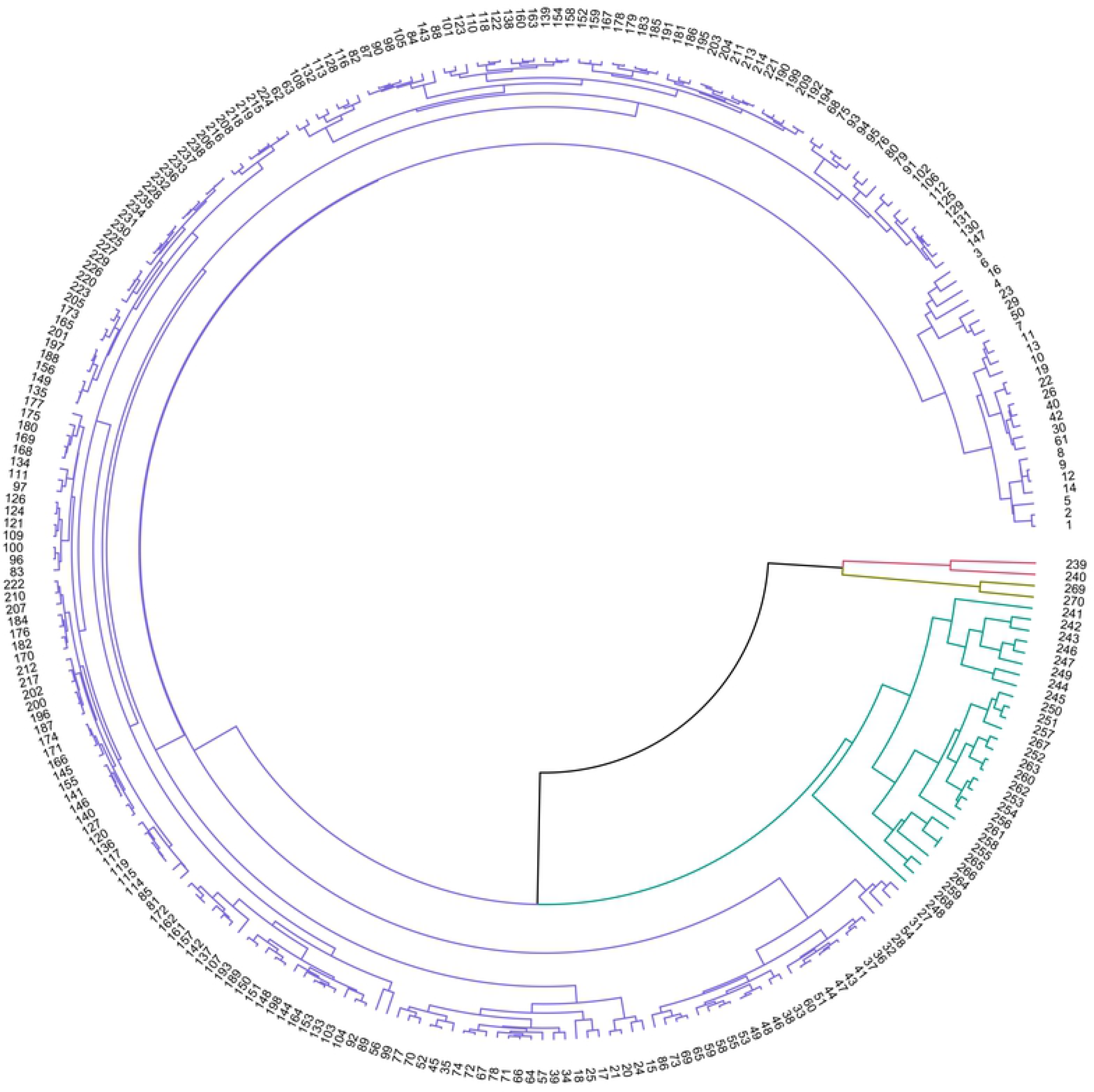
Cluster Diagram obtained from Agglomerative hierarchical clustering using Euclidean Distance as matric function. Four clusters were formed at 3.9 Clustering scale. Cluster 1 (Howl), Cluster 2 (Whimper), Cluster 3 (Social Squeak) and Cluster 4 (Whine) are denoted by Blue, Red, Green and Brown Respectively.

**Figure 4.**
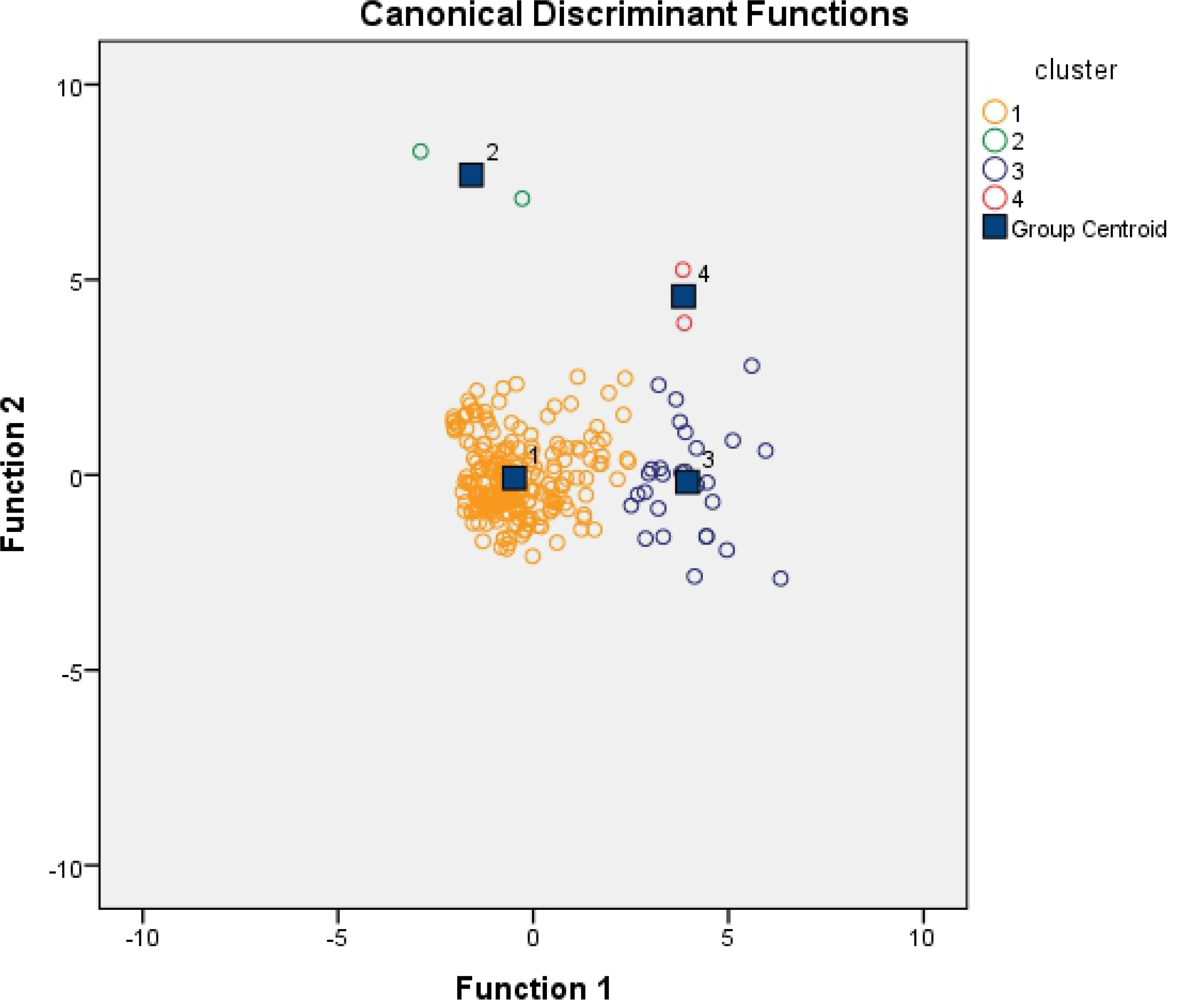
The plot of Discriminant Function Analysis (DFA) using PCA value for 270 vocalisation data from Indian Wolf. Different colours represent different call type.

### Discriminant Function Analysis (DFA)

DFA achieved 95.9 % accuracy of vocal group identification using two PCA values (Table 3). Each of the four groups has a distinct group centroids. Graphical representation using two discriminant functions (DF1 and DF2) shows that vocal clusters do not overlap (Figure 5).

**Table 3.** Classification Results of Discriminant Function Analysis (DFA). 95.9% Vocal clusters (estimated from Agglomerative hierarchical clustering) are identified correctly.

|  |  | cluster | Predicted Group Membership |  |  |  | Total |
| --- | --- | --- | --- | --- | --- | --- | --- |
|  |  |  | 1 | 2 | 3 | 4 |  |
| Predicted in Cluster analysis | Count | 1 | 229 | 0 | 8 | 1 | 238 |
|  |  | 2 | 0 | 2 | 0 | 0 | 2 |
|  |  | 3 | 0 | 0 | 26 | 2 | 28 |
|  |  | 4 | 0 | 0 | 0 | 2 | 2 |
|  | % Correct | 1 | 96.2 | .0 | 3.4 | .4 | 100.0 |
|  |  | 2 | .0 | 100.0 | .0 | .0 | 100.0 |
|  |  | 3 | .0 | .0 | 92.9 | 7.1 | 100.0 |
|  |  | 4 | .0 | .0 | .0 | 100.0 | 100.0 |

**Figure 5.**
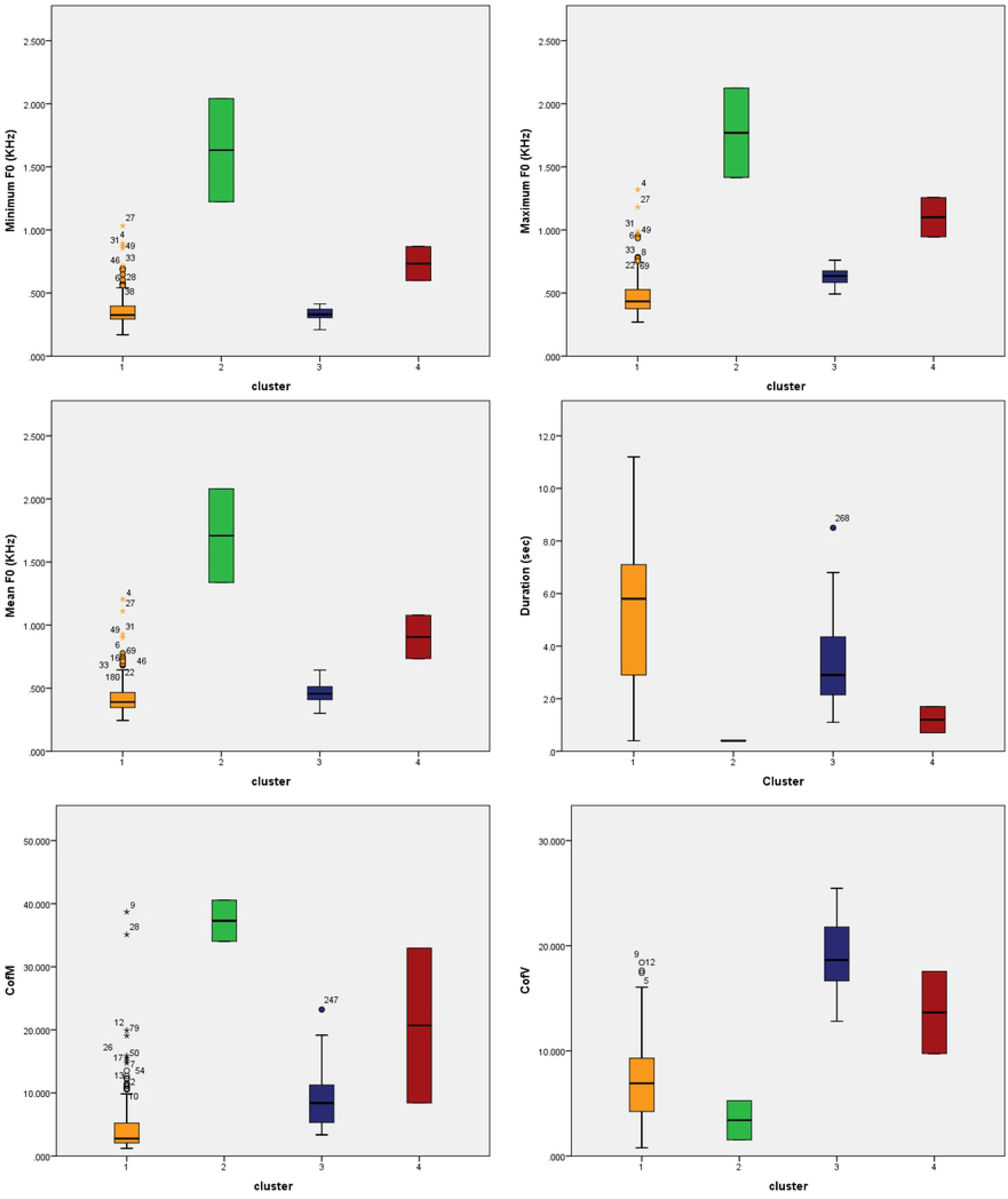
Box plot of variation among acoustic variables between different call type. a. Variation among minimum frequency, b. Variation among maximum frequency, c. Variation among Mean frequency, d. Variation among duration of the call, e. Variation among coefficient of frequency modulation, f. variation among coefficient of frequency variation.

### Box plot using a simple scalar variable

The whisker box plot represents the variation among the acoustic variables within the four identified call types (Figure 6). Call type 1 had the longest duration (5.214±2.49 Sec) whereas call type 2 showed the shortest duration among the four recognised groups (0.4±0) (N=2). Type 2 calls are also high frequency modulated (37.296±4.601) (variation in frequency per unit time). However, frequency variation (around the mean) is highest in type 3 calls (18.778±3.587) (Table 4).

**Table 4.**
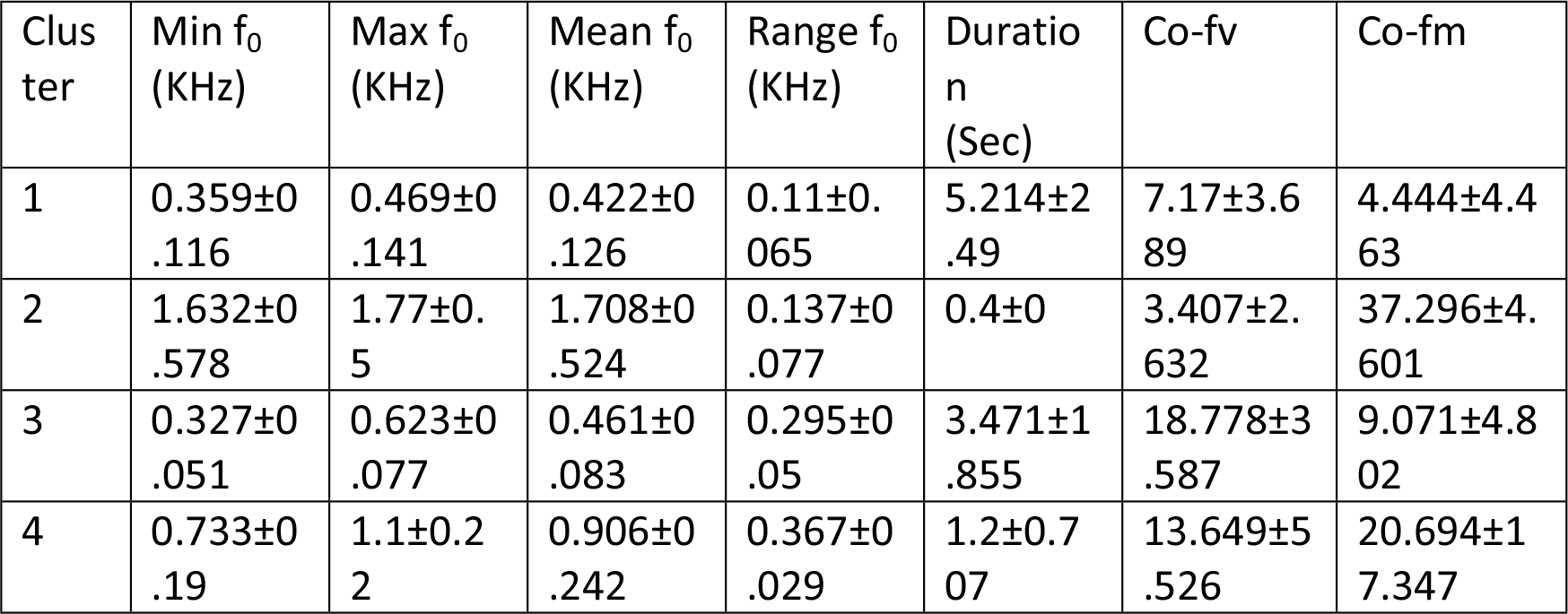
Variation among important acoustic variables within the four identified call types.

**Figure 6.**
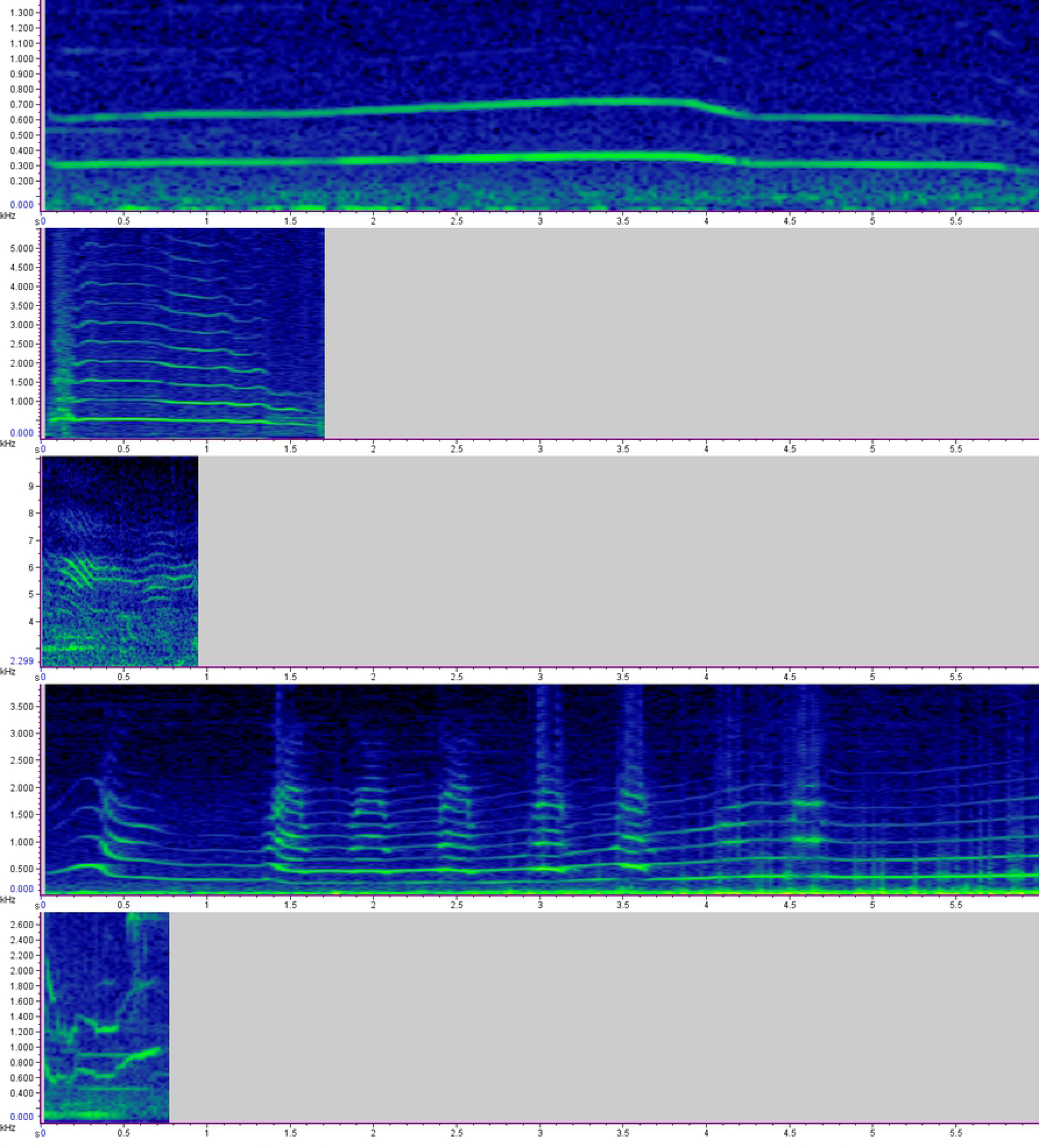
Spectrograms of different type of vocalisation in Indian wolf. a. Howl (type 1); b. Bark-Howl (type 1 subtype); c. Whimper (type 2), d. Social Squeak (type 3); e. Whine (Type 4);

## Discussion

This study provides a quantitative assessment of the vocalisations of the Indian wolf subspecies. Our results show that there are four statistically classified groups of Indian wolf vocalisations based on ten captive individuals and nine free-ranging Indian wolf packs. Though the Four to Six solution groups showed a narrow difference in their average silhouette values based on *silhouette plot* analysis, the four cluster solution was found to be the most significant based from the global maxima. This characterisation of vocalisations provides a first step to evaluating the function and contextual use of different types of vocalisations in these canids.

The first call type is identified as a howl (Figure 7a), which is the most dominant and prolonged (5.214±2.49 sec) call type in our dataset. The fundamental frequency of howl ranged from 0.359 KHz (±0.116) minimum to 0.469 KHz (±0.141) maximum (N=238). Besides having smaller body size, the mean Fundamental Frequency of the Indian wolf howl (0.422±0.126 KHz) was similar as compared to other Holarctic clade subspecies reported by Hennelly et al. [46]. Since the howl is the most detectable vocalisation used in long-range social cohesion and territorial advertisement [8,22], our high howl sample size shouldn’t be considered as the most dominant vocalisation. Barking-howl, which was mentioned by many authors as a common type of mix vocalisation in wolves [22,27,47], falls under the same cluster along with howling (Figure 7b). From our field observations, wolves bark in defence to an immediate threat. In one such occasion, the she-wolf of a pack started barking at nearby villagers to protect her cubs and did not stop until all the three pups ran away to a safer distance from the villagers.

The second call type has the highest frequency modulation (37.296±4.601 KHz) and is commonly known as a whimper (Figure 7c). The whimper is low intensity but high-pitched sound and unlike howls, it is used for short distance communication among pack members. [20,22] (mean fundamental frequency = 1.708±0.524 KHz). This short duration (0.4±0 KHz) vocalisation is reported to be associated with submissive or friendly greeting behaviour [22,47]. Since it is not audible from more than one to two hundred yards away [47], our dataset contains only a few observations (difference sources) of this type of call (N=2). While our study provides some initial insight into the acoustic structure of this vocalization, further sampling will be needed to robustly characterize the acoustic structure of the whimper.

The third group of Indian wolf vocalisations can be termed as ‘*social squeak*’ (Figure 7c), following observations by previous studies (Mech 1981, Crisler 1959, Fentress 1967). This high-frequency variable vocalisation (18.778±3.587 KHz) in the Indian wolf is similar to ‘talking’ as termed by Crisler [48]. Social Squeak has a minimum frequency of 0.327 KHz (±0.051) to Maximum of 0.623 KHz (±0.077) (N=28).

Lastly, our fourth group we identified as the whine (Figure 7e), which is characterized as a short duration vocalisation (1.2±0.707 Sec). The whine in the Indian wolf (*Canis lupus pallipes*) is longer than the whine reported in Italian wolf (0.13±0.10 Sec) [27]. Although our data set is small (N=2; from two different individuals) to interpret robustly, the mean fundamental frequency of Indian wolf whine (0.906±0.242 KHz) has a similar frequency as the Italian wolf (0.979±0.109 KHz) [27].

Overall, our study identified four statistically classified groups of vocalisations of the Indian wolf. While the howl has been extensively studied in its behavioural function and variation across subspecies [8,46], less is known about the short-range communication among wolves. For example, our study identified the ‘Social Squeak’, or the ‘talking’ like vocalisation in the wolf, yet little is known about its function in wolf packs and if it’s a common communication across different canid species and within domestic dogs. Further research on a larger dataset of Indian wolf vocalisations can develop a more robust classification of the vocal repertoire and can be used for future studies on the contextual use and variations across different wolf subspecies. More broadly, the species within the *Canis* clade vary in their body sizes, social structure, and habitats. Describing the vocal repertoires of various canid species can give insight into the factors influencing vocal communication within the genus *Canis*.

## Acknowledgements

We want to thank the authority and staffs of the Jaipur Zoo and Maharashtra Forest Department for permitting to conduct this research. We sincerely acknowledge the funding agencies, Department of Science and Technology, India and Forest Department of Maharashtra. We appreciate all our field personals and wildlife enthusiast groups (Pune Wolfgang, Mihir Godbole, Vineet Arora, R. V. Kasar, Rajesh Pardeshi, Sawan Behkar and others) from Maharashtra who helped in local information gathering and various logistic arrangement during data collection. The first author is grateful to Holly-Root Gutteridge, who helped me in learning feature extraction during the initial days. We acknowledge the effort of Shivam Shrotriya and Nilanjan Chatterjee for assistance during the analysis. We would like to show our gratitude to our friends and colleagues whose insight and comments considerably improved the manuscript. We are delighted for having the continuous support and facility from Wildlife Institute of India, Dehradun.

## Supporting Information

**S1 File. The Variables, Clusters and Other Details information of every calls.**

